# Temperate gut phages are prevalent, diverse, and predominantly inactive

**DOI:** 10.1101/2023.08.17.553642

**Authors:** Sofia Dahlman, Laura Avellaneda-Franco, Ciaran Kett, Dinesh Subedi, Remy B. Young, Jodee A. Gould, Emily L. Rutten, Emily L. Gulliver, Christopher J.R. Turkington, Neda Nezam-Abadi, Juris A. Grasis, Dena Lyras, Robert A. Edwards, Samuel C. Forster, Jeremy J. Barr

**Affiliations:** School of Biological Sciences, Monash University, Clayton, VIC, 3800, Australia; Centre for Innate Immunity and Infectious Disease, Hudson Institute of Medical Research, Melbourne, 3168, Australia; Department of Molecular and Translational Sciences, Monash University, Clayton, Victoria, 3800, Australia; APC Microbiome Ireland & School of Microbiology, University College Cork, Cork, Ireland; Department of Molecular and Cell Biology, University of California, Merced, CA 95343, USA; Monash Biomedicine Discovery Institute Department of Microbiology, Monash University, VIC, 3800, Australia; College of Science and Engineering, Flinders University, Bedford Park, SA 5042, Australia

**Author notes:** These authors contributed equally to this work (Samuel C. Forster, Jeremy J. Barr).

## Abstract

Large-scale metagenomic and data mining efforts have uncovered an expansive diversity of bacteriophages (phages) within the human gut^1–3^. These insights include broader phage populational dynamics such as temporal stability^4^, interindividual uniqueness^5,6^ and potential associations to specific disease states^7,8^. However, the functional understanding of phage-host interactions and their impacts within this complex ecosystem have been limited due to a lack of cultured isolates for experimental validation. Here we characterise 125 active prophages originating from 252 diverse human gut bacterial isolates using seven different induction conditions to substantially expand the experimentally validated temperate phage-host pairs originating from the human gut. Importantly, only 17% of computationally predicted prophages were induced with common induction agents and these exhibited distinct gene patterns compared to non-induced predictions. Active Bacteroidota prophages were among the most prevalent members of the gut virome, with extensive use of diversity generating retroelements and exhibiting broad host ranges. Moreover, active polylysogeny was present in 52% of studied gut lysogens and led to coordinated prophage induction across diverse conditions. This study represents a substantial expansion of experimentally validated gut prophages, providing key insights into their diversity and genetics, including a genetic pathway for prophage domestication and demonstration that differential induction was complex and influenced by divergent prophage integration sites. More broadly, it highlights the importance of experimental validation alongside genomic based computational prediction to enable further functional understanding of these commensal viruses within the human gut.

## Main

The human gut microbiota consists of a plethora of microorganisms along with the viruses that infect them, including phages. These viruses are thought to shape the gut microbial community through predation, horizontal gene transfer, and lysogenic conversion^9,10^. Recent advances in computational mining of gut metagenomes have revealed an expansive collection of viral metagenome-assembled genomes (vMAGs) and efforts into cataloguing this diversity have led to the discovery of several important viral families^1–3,6,11,12^. In addition, lysogeny is common within the gut with up to 90% of bacteria predicted to harbour prophages^13,14^. However, the extent to which these prophages re-enter lytic replication remains unknown. For example, the inactivation of resident prophages represents a common strategy whereby the bacterial population can escape lysis while maintaining beneficial temperate phage genes^15,16^. Further, initiation of lytic replication by resident prophages is complex, involving both host and phage specific cues^17,18^. Within the gut, little is known about temperate phages and how they interact with commensals.

### Induction of prophages from human gut isolates

Advances in cultivation of the microbiota have enabled the isolation and archiving of previously ‘unculturable’ gut bacterial species^19,20^, along with their phages. Here we utilise a collection of 252 human gut bacterial isolates (50 Actinomycetota, 1 Fusobacteriota, 51 Bacillota, 57 Pseudomonadota and 93 Bacteroidota) to computationally identify and experimentally validate inducible prophages (Supplementary Table 1). Applying seven induction conditions (0.3 and 3 μg/mL Mitomycin C, 0.5 mM Hydrogen peroxide, 3.7 mg/mL and 37 mg/mL Stevia, 50% Carbon depletion, 100% short-chain fatty acid (SCFA) depletion) we recovered and sequenced 431 viral induction samples passing our filtering criteria (Extended Data Fig. 1, Supplementary Table 2) resulting in the detection of 125 inducible gut prophages representing 63 phage species (95% ANI over 85% alignment fraction, AF)^21^ from 73 bacterial isolates (5 Actinomycetota, 1 Fusobacteriota, 10 Bacillota, 17 Pseudomonadota and 40 Bacteroidota)(Fig. 1)

**Fig. 1.**
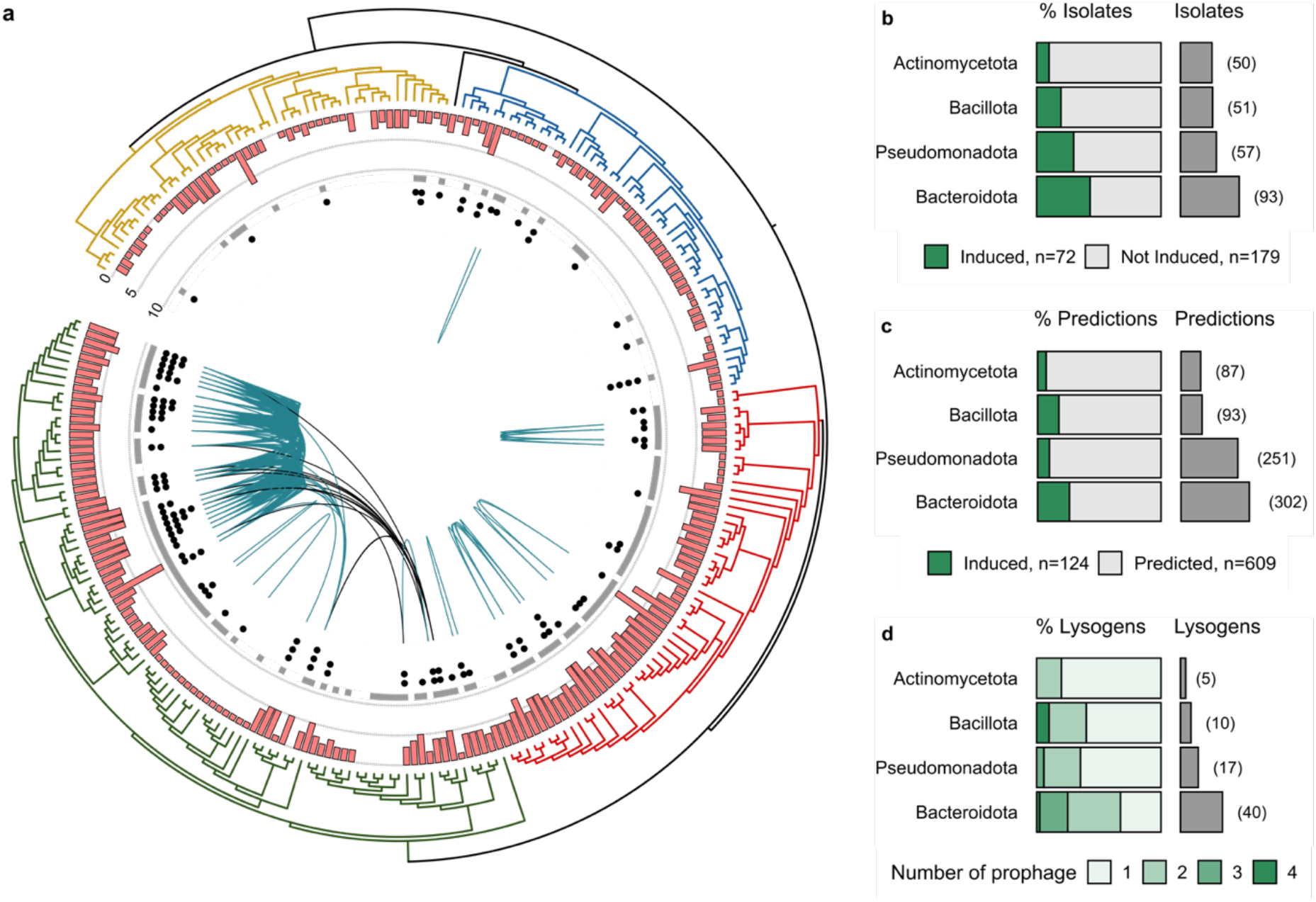
Distribution of induced prophages within gut bacterial isolates. **a**, Phylogenetic tree of gut bacterial isolates used for induction. Actinomycetota in yellow (n=50), Fusobacteriota in black (n=1), Bacillota in blue (n=51), Pseudomonadota in red (n=57) and Bacteroidota in green (n=93). Pink bars represent number of high quality (> 50% completeness) predicted prophage regions, inner ring shows whether the isolate was sequenced in at least one phage induction sample (grey), dots represent induced prophages within each isolate (black). Green lines connect bacterial isolates sharing the same induced phage and black lines connect isolates of different genera harbouring the same prophage species. **b**, Percentage induced and total number isolates. **c**, Percentage induced and total number high quality prophage predictions. **d**, Distribution of induced single and polylysogen per bacterial phyla. Fusobacteriota excluded from **b, c** and **d** as single isolate.

### Only a fraction of predicted gut prophages were inducible

Consistent with previous reports of substantial lysogeny within the human gut^13^, 237 isolates (94%) were computationally predicted to contain high-quality prophage regions (Fig. 1a). However, only 29% (73/252) of isolates were induced under the conditions investigated and 17% (125/736) of the high-quality predictions, or 23% (63/274) of high-quality prophage species, corresponded to an experimentally inducible prophage (Fig 1b-c). Moreover, the fraction of inducible to predicted prophages reported here coincides with recent reports from gut metagenome sequencing (20-36%), that indicate only a minority of gut prophages readily undergo lytic replication^22,23^. While our experimental approach does not provide comprehensive identification of all gut prophage due to factors including detection limits and absence of specific stimuli, it is likely that most predictions within our dataset represent inactive gut prophage; referred to as cryptic prophages^16^.

### Bacteroidota isolates exhibit a greater proportion of active prophage

The highest concordance between active and predicted prophage regions was observed within Bacteroidota isolates, where 78 predictions (26%) from 40 isolates (43%) were inducible (Fig 1b-c). Comparatively in Pseudomonadota, which harboured the highest number of predicted prophages (4.5 per isolate), just 10% of prophages were found to be active. Polylysogeny was most prevalent within the Bacteroidota isolates where 27/40 (68%) of lysogens harboured more than one active prophage (Fig. 1d), compared to 11/33 isolates (33%) across the other phyla (*p*=0.005; Fisher exact Test). While most phage species infected a single host, seven out of 27 Bacteroidota prophages were found actively replicating across bacterial species, three of which were found to be induced across bacterial isolates from different genera (Fig. 1a).

### Temperate phage taxonomy in inducible gut isolates

Given the inherent challenges in assigning taxonomy to phages^24^, we applied both a gene voting based search and a gene sharing network method using vContact2 (Fig. 2a)^25^ with taxonomy assigned by highest shared taxonomic resolution between methods. The resulting classification assigned 124 phages to the Caudoviricetes order and the other within the Faserviricetes order. In total, 27% (34/125) of phage could be assigned to ICTV accepted taxa at family level or lower. These belonged to previously reported phage taxa infecting Pseudomonadota (Bcepmuvirus, Glaedevirus, Punavirus, Uetakevirus and Peduoviridae), one Spbetavirus infecting Bacillota and 16 prophages belonging to the Winoviridae family infecting Bacteroidota. Although lacking ICTV classification, 29 genomes could be grouped into viral clusters (approximately genus level) together with previously described phages (Supplementary Table 3). Notably, 19 of these clustered with Hankyphage, a recently described virus thought to lysogenize several Bacteroides species^26^. Further taxonomic classification grouped ten prophages at the species level with Hankyphage whereas the remaining nine clustered into seven putative novel species, forming a putative novel genus which we name “Hankyvirus” after the original phage characterised (Extended Data Fig. 2a). Comparing the Hankyvirus species to bacterial genomes in NCBI RefSeq database (95% ANI over 85% AF), we identified 52 host species originating from nine genera and five families, indicating a broad host range of this genus (Extended Data Fig. 2b). Correspondingly, we find two Hankyvirus species induced within both *Bacteroides* and *Phocaeicola* isolates within our set, providing experimental validation of these phages as actively replicating across these two host genera.

**Fig.2.**
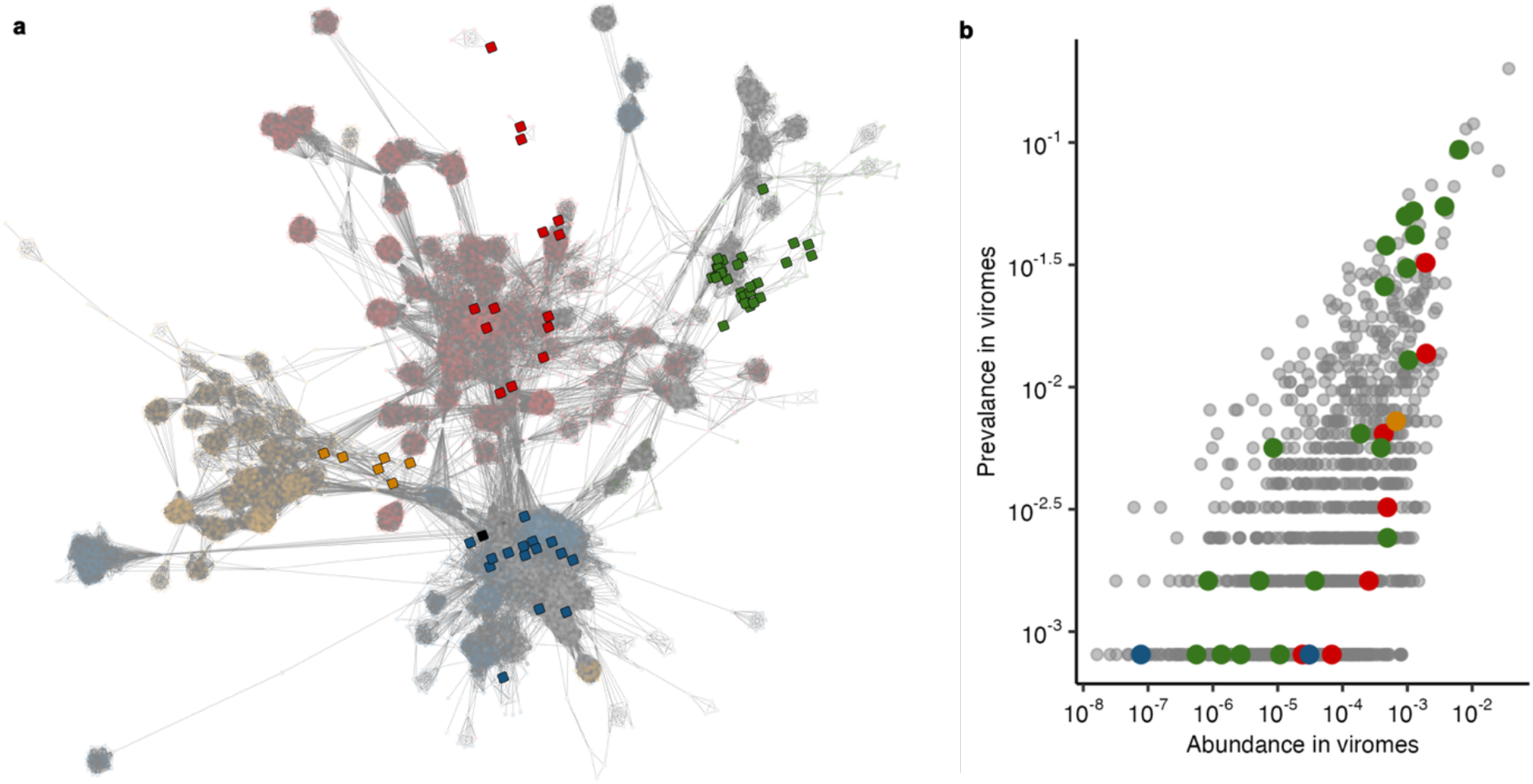
Taxonomy and prevalence of induced temperate phages within gut viromes. **a**, Gene sharing network of temperate induced phage species (solid squares, n=63) coloured by host; Actinomycetota in yellow (n=6), Fusobacteriota in black (n=1), Bacillota in blue (n=15), Pseudomonadota in red (n=14) and Bacteroidota in green (n=27). Database representatives (n=9920) in translucence and coloured by host phyla when applicable, otherwise in grey. **b**, Mean fractional abundance and detection frequency (prevalence) of Caudoviricetes phages within 1232 viromes originating from the human gut. A minimum of 70% coverage over the length of the phage was required to be counted as present within a virome. The bacterial host phyla of induced temperate phage species coloured as in **a** whereas database reference genomes (n=1085) are shown in grey.

### Inducible temperate phages are prevalent within gut viromes

We next sought to understand the prevalence of our inducible prophages within human gut viromes. Approximately half of the inducible prophage species (31/63) could be detected in gut viromes (n=1232, Fig. 2b, Supplementary Table 4). LoVEphage, a recently discovered Bacteroidota phage^11,12^, was most common, with detection in ∼9% (116/1232) of the viromes and representing up to 89% of reads within some viromes. Three phages in our collection were species-level members of LoVEphage, induced from *Bacteroides thetaiotaomicron, Phocaeicola dorei* and *Phocaeicola vulgatus* hosts (Extended Data Fig. 2b). An additional seven temperate phage species were detected in 3-5% of gut viromes (Supplementary Table 2). These included the prototypical Hankyphage and two additional Hankyvirus species, three previously uncharacterised Bacteroidota phages and a Uetakevirus infecting *Escherichia coli*.

### DGRs are common within Bacteroidota gut prophages

Discernible integrase or site-specific recombination genes, both of which are used as hallmark genes for a temperate lifestyle^4,5^ was absent in 30% (19/63) of our inducible phage species, including Hankyviruses. We found transposases in 11 of these viruses, while the remaining eight lacked any discernible integration genes, illustrating the difficulty in assigning phage lifestyle based on genomic data alone. Viral diversity generating retroelements (DGRs) are prevalent within the gut virome and tail-targeting DGRs have been shown to enable rapid host switching in a *Bordetella* phage^27,28^. We found DGRs in 20% (13/63) of inducible phage species, the majority of which were seen in Bacteroidota phages, where 44% (12/27) of species encoded DGRs targeting known and genomically predicted tail proteins. Notably, four of these species encoded a second variable region (VR) targeting genes distal from the reverse transcriptase (RT) cassette (Extended Data Fig. 3)^11^. The second VR was found in proximity to counter defence genes, such as DNA methyltransferase, indicating a possible involvement of DGRs in the phage-host arms race reaching beyond host range expansion through tail fiber diversification^26^.

### Cryptic prophages are enriched with accessory genes

The presence of cryptic (inactive) prophages is known to provide their host with adaptive fitness advantages^16^. A bimodal length distribution of the predicted prophage regions across all host phyla within our collection was observed (Extended Data Fig. 4a). This bimodal size distribution of prophage genomes is thought to result from an active retention of smaller prophage-like elements and an ongoing influx of new complete prophages^15^. When grouped by completeness, our induced prophages clustered with the larger peak corresponding to high-quality predictions (>50% complete, n=736), whereas the smaller peak corresponded to sequences with low completeness scores (<50% complete, n=1236) (Fig. 3a). To investigate whether there were differences in gene content between these groups, we performed gene enrichment analysis of annotated PHROG gene categories^29^. Small prophage genomes lacked essential phage genes (such as structural, head and packaging, and lysis genes) but were enriched in accessory and genes with unknown function (Extended Data Fig. 4b). Comparing induced prophages to high-quality predictions, the induced prophages showed enrichment in total gene frequency for head associated genes (*p*=7.3×10^−5^, Fisher Exact Test) and enrichment in presence-absence frequency in genes essential for phage function including packaging (*p*=4.2×10^−10^, Fisher Exact Test), connector (*p*=5×10^−2^, Fisher Exact Test) and lysis (*p*=1×10^−3^, Fisher Exact Test). In contrast, high-quality prophage predictions showed enrichment in total gene frequency for accessory (*p*=8.7×10^−4^, Fisher Exact Test) and unknown genes (*p*=5.3×10^−3^, Fisher Exact Test)(Fig. 3b). Next, we sought to investigate potential genetic mechanisms leading to prophage inactivation, by identifying cryptic (non-induced) prophages aligning over at least 85% the length of an inducible prophage within our sample set. To classify these prophages as cryptic, rather than induced below our limit of detection, we restricted the cryptic prophage set to those that had been sequenced (and not induced) in the same condition(s) as their inducible counterparts. This resulted in a total of 211 active-cryptic prophage pairs, between 57 active and 48 cryptic prophages. No significant changes were found in gene frequency (*p*>0.05, Fisher exact test) indicating that while gene loss may be characteristic of cryptic prophages, it is unlikely to be the initial cause of inactivation. Further, while we detected 65 homologous gene transfer and 9 insertion/deletion events within the active-cryptic pairs, there was no significant difference in the number of events when compared to a set of high sequence similarity active-active pairs (n=205, *p*=0.28 and *p*=0.62, Fisher exact test). Comparing host ANI between the active-active and active-cryptic prophage pairs (Extended Data Fig. 4c) we found no association between host ANI and prophage induction (*p*=0.13, Pearsons correlation, Extended Data Fig. 4e), suggesting that prophage inactivation is not driven by integration into divergent non-permissive hosts or increased diversification of cryptic prophage hosts.

**Fig. 3.**
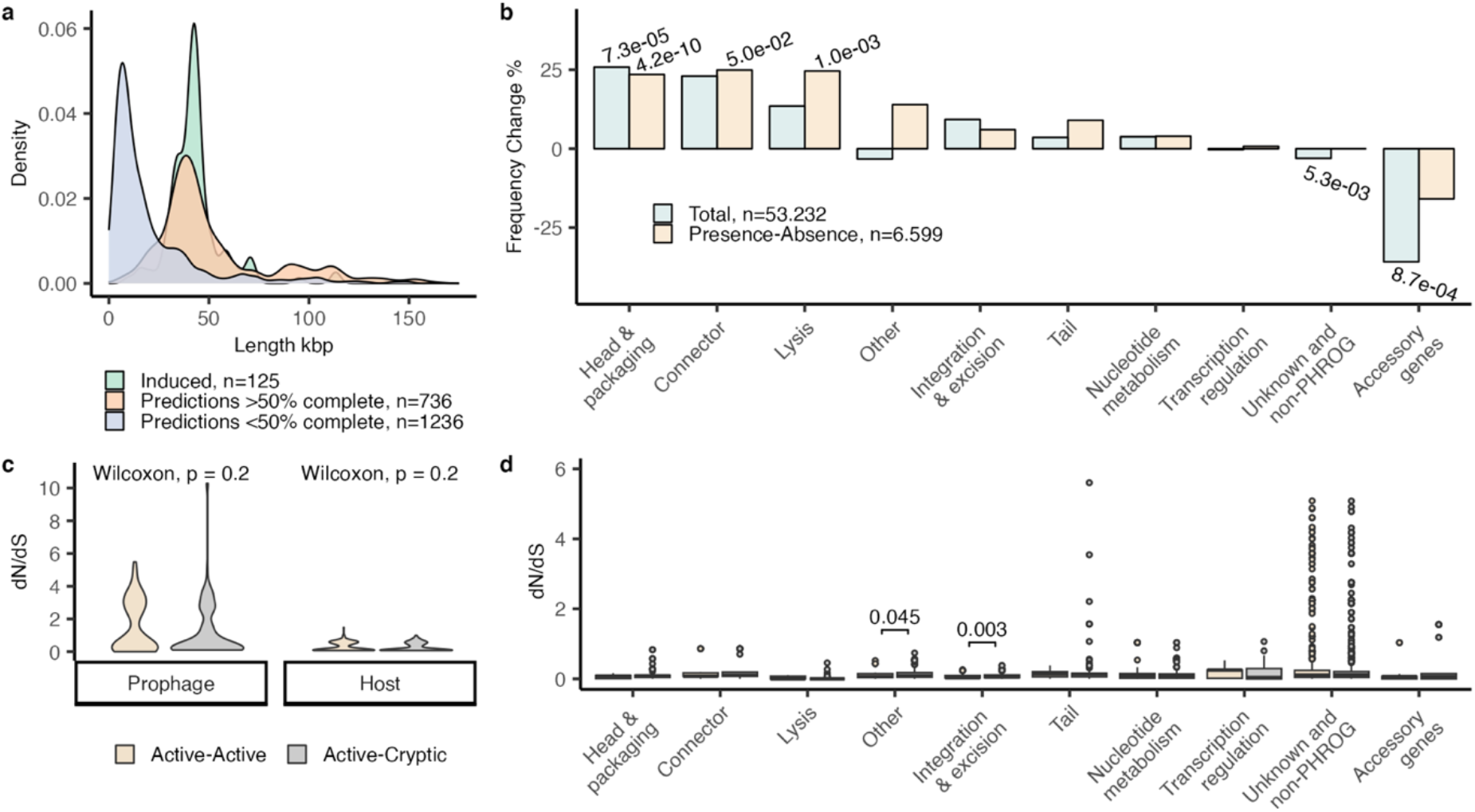
Comparison of induced versus predicted prophages. **a**, Length distribution of induced, high-quality prediction (>50% complete) and low-quality prediction (<50% complete) prophage genomes. **b**, Percentage frequency change in PHROG gene categories between induced and high-quality (>50% complete) prediction prophage genomes, counting total genes (blue) or presence of at least one gene per category and genome (yellow). Significant *p* values calculated using Fisher’s exact test and adjusted by Hochberg method shown above bars. **c**, dN/dS rates between active-active or active-cryptic prophage pairs (left) and between their hosts (right). Wilcoxon signed-rank test showed no significance. **d**, dN/dS rates of PHROG gene categories between active-active and active-cryptic prophage pairs. Significant *p* values calculated using Wilcoxon test and adjusted by Hochberg method shown above brackets.

### Cryptic prophages have elevated mutation rates in integration and excision genes

To investigate whether cryptic prophages harbor elevated number of muations, we measured the ratio of non-synonomous to synonomous substitution rates (dN/dS) within the set of active and cryptic prophage pairs, and their associated hosts. We found an overall elevated mutation rate in prophages (mean=1.4/median=0.55) compared to host genome (mean=0.3/median=0.13, *p*<2.2e-16, Wilcoxon test), but no significant difference between active or cryptic prophage pairs or their host genomes (*p*=0.2, Wilcoxon test, Fig. 3c). Comparing gene substitution rates, we find 80 genes, (63 of which where unknown genes), with elevated dN/dS rates (>1) indicating positive or diversifying selection, with 48% (38/80) of these genes being associated with DGR’s (Fig. 3d). Consistent with inactivity, a significant increase in the dN/dS substitution rate within cryptic prophage genes involved in integration and excision (*p*=0.003, Wilcoxon test) was observed.

### Phyla-specific cues may govern prophage induction within gut isolates

To understand the factors contributing to prophage induction within our tested isolates we compared induced prophages across the seven induction conditions. Combined, the two concentrations of Mitomycin C induced the largest number of prophages (n=69), induced the most Pseudomonadota (n=17), and was the only condition in which Actinomycetota prophages were induced (n=7) (Fig. 4a). Hydrogen peroxide induced 43 prophages including the largest number of Bacteroidota prophages (n=35). However, these well-known induction agents exhibited only a marginally increased induction rate compared to spontaneous induction during standard growth condition (n=36). Considerable overlap was observed between prophage induction in standard media and induction agents (Mitomycin C; n= 25, Hydrogen peroxide; n=15, Stevia; n=19, Carbon depletion; n=9 and SCFA depletion; n=11). As such, whereas certain agents promoted induction within specific phyla, induction could not exclusively be attributed to their action.

**Fig. 4.**
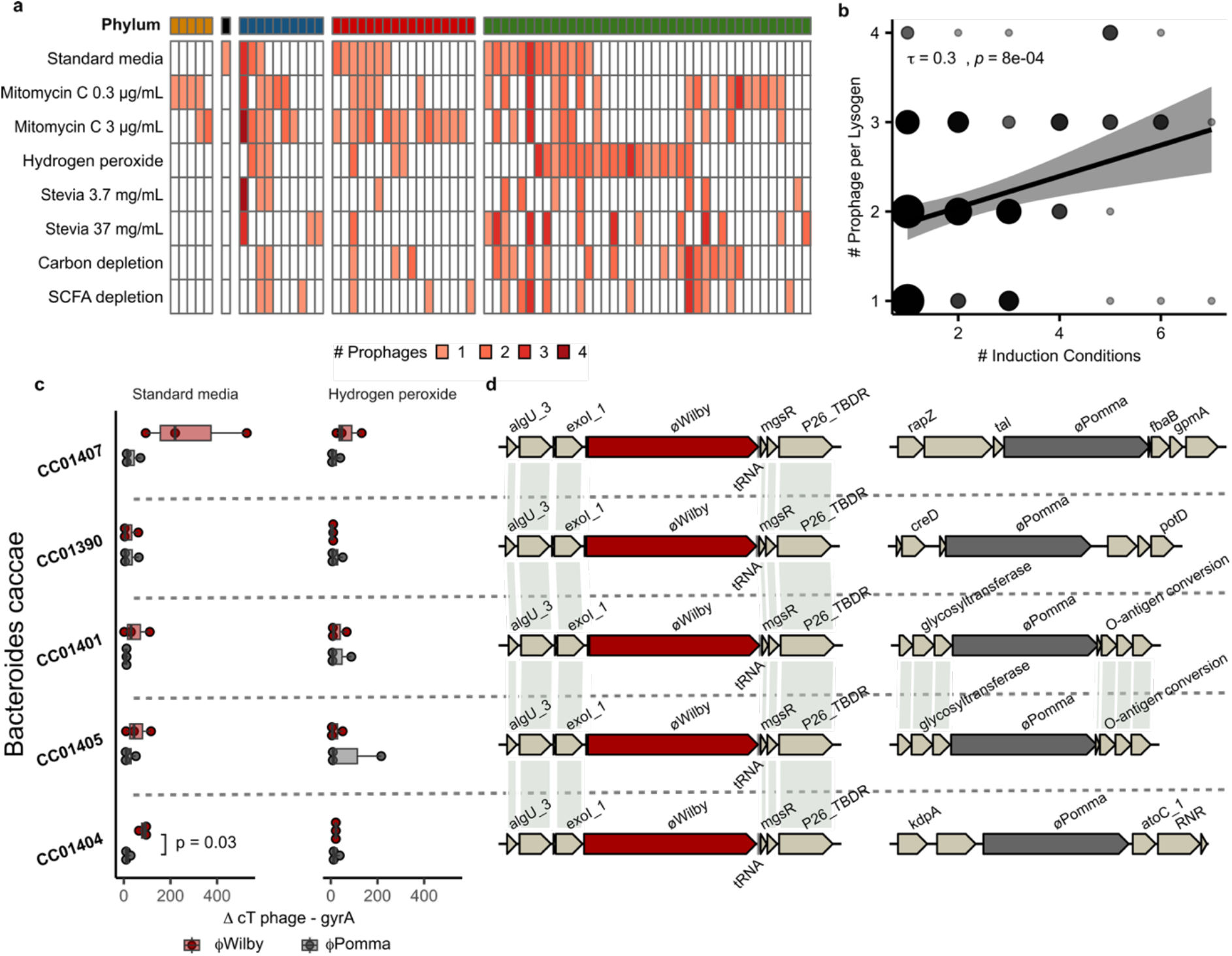
Comparison of induction agents and analysis of polylysogeny within gut isolates. **a**, Number induced prophages per samples (condition in rows and isolates in columns). Isolate phylum show in in top bar; Actinomycetota in yellow, Fusobacteriota in black, Bacillota in blue, Pseudomonadota in red, and Bacteroidota in green. **b**, Kendall’s rank correlation between number of prophages within lysogens and number of conditions in which phages were detected as induced (size based on number observations). **c**, Fold change induced prophage over background in isolates grown in only Standard media only (left, *n*=3) or with addition of Hydrogen peroxide (right, *n*=3). Significant *p* values calculated using paired t-test shown above brackets. **d**, Genome location of prophage ΦWilby (red) and ΦPomma (grey). Lines connect genes with 100% AAI.

### Polylysogenic prophage genomic location influences induction in near identical isolates

Next, we investigated the influence of polylysogeny on induction, observing a positive correlation between the number of co-inhabiting active prophages and conditions leading to induction (*τ*=0.3, *p*=8e-04, Kendall’s rank correlation, Fig. 4b). Prophages residing in polylysogens (n=90) were induced on average in 2.4 conditions compared to 1.9 conditions in single lysogens (n=35, *p*=0.02, Wilcoxon test). This suggests polylysogeny may promote simultaneous prophage induction across diverse conditions and reduce stability within lysogens. To investigate differential induction within polylysogens, we measured the abundance of phage DNA in supernatants of five highly similar (99% ANI) *Bacteroidota caccae* isolates harbouring the same two prophages (ΦWilby and ΦPomma). We identified preferential induction of ΦWilby within standard media (*p*=0.026), but not in Hydrogen peroxide treated samples (*p=*0.9, Wilcoxon test), with isolate CC01404 demonstrating the most marked difference (*p*=0.03, paired t-test, Fig. 4c). Calculating the ratio of ΦWilby over ΦPomma within each isolate, we found a significant variance of means between the isolates in both standard media (*p*=0.013) and Hydrogen peroxide (*p*=0.0002, ANOVA). These results implied host genetic background, even within highly similar isolates, may affect prophage induction. We previously identified phage ΦPomma as a transposable prophage, which does not utilise site specific integration, but randomly inserts into the host genome^30^. To investigate the prophage integration sites within our isolates, we undertook long read sequencing on the five *B. caccae* strains. Genomic analysis identified ΦWilby integrated into the same tRNA gene location, which is characteristic of site-specific integration; however, the transposable prophage ΦPomma was found in four different genomic locations within the five isolates (Fig. 4d) implicating integration site as the primary driver for the observed differential induction in these isolates.

## Discussion

The high microbial load within the human gut represents an optimal environment for temperate phages, as frequent interactions with their hosts provide ample opportunity for lysogeny^14^. Concordantly, the majority of bacteria within the gut are predicted to be lysogens, with up to 90% of bacteria harbouring at least one prophage^13,14^. However, whether these predictions represent active prophages able to re-enter the lytic lifecycle, or cryptic prophages trapped in the bacterial chromosome, is less well understood. Leveraging our defined culture collection of 252 gut bacterial isolates, we also predict the majority to harbour prophage-like elements (94%), but find that only a fraction of predicted prophages were capable of active replication (17%). Considering little is known about prophage triggers within the gut, it is plausible that some of our isolates carry active prophages that were not induced in this study. However, host domestication of prophages is common, as the carriage of active prophages comes with added fitness costs^15^. We propose that whereas the genetic pool of integrated prophages within the gut is large, only a fraction of these will readily re-enter the lytic lifecycle. The genetic mechanism behind the widespread inactivation of prophages observed within gut isolates remain largely unknown. In this study, we detect distinct gene enrichment patterns where cryptic prophages encoded fewer structural and lysis-associate genes and had higher non-synonymous substitution rates in integrase and excision-related genes, providing a genetic pathway towards the inactivation and domestication of gut prophages.

A considerable portion of our isolates (52%) with inducible prophages were polylysogens, harbouring more than one replicating prophage^31^. We found a positive correlation between polylysogeny and number of successful induction conditions. These findings are consistent with previous reports of polylysogenic *Salmonella* strains where phage antirepressor proteins can inactivate repressors made by hetero-immune prophages, thereby synchronising prophage induction^32^. Finally, we provide evidence for phyla-specific induction cues and show that differential induction of polylysogenic prophages varied between near identical isolates due to divergent prophage integration sites within the host genome. Thus, differential prophage induction was influenced by the inducing agent, host phyla, polylysogeny, and prophage integration site. In conclusion, we demonstrate the feasibility of culture-based approaches to provide new insights into phage-host interaction within human-associated commensals.

## Acknowledgements

This work was supported by the Australian Discovery Project grant (DP210103296). In addition, S.D and L.A.F were supported by Monash University Postgraduate Research Scholarship funding their doctoral studies and S.C.F. is supported by a CSL Centenary Fellowship. The authors would also like to acknowledge the Monash eResearch Team and the support of the Victorian State Government Operational Infrastructure Scheme.

## Author contributions

J.J.B. and S.C.F. conceived and designed the study. R.A.E., D.L. and J.A.G. contributed ideas and expertise. S.D. performed *in vitro* inductions, molecular work, sequencing, informatic analyses, and most of the data preparation. L.A. assisted with informatic analyses, data interpretation, molecular work and sequencing. C.K. performed DNA extractions. D.S. performed qPCR assays. R.B.Y, J.A.G., and E.R. assisted with molecular work and sequencing. E.R., E.L.G and R.B.Y isolated and assisted with the cultured bacterial isolates. C.J.R.T. and N.N.A. assisted with identification of induced prophages. All authors reviewed and discussed the manuscript. S.D., S.C.F. and J.J.B. wrote the paper. J.J.B. supervised all aspects of the work and all authors approved the final manuscript

## Data Availability

All data are available from the corresponding author upon reasonable request

