## Supplementary materials and methods for "Temperate gut phages are prevalent, diverse, and predominantly inactive"

### Bacterial culture conditions

A culture collection of 252 bacterial isolates previously isolated and sequenced from human gut samples were used for phage induction. All bacterial culture work was performed in yeast-extract casitone fatty acid (YCFA) media at 37 °C in anerobic conditions (Whitely A95 anaerobic workstation) containing 10% carbon dioxide, 10% hydrogen and 80% nitrogen[1]. Each isolate was streaked onto YCFA agar plates and incubated for 24 hours before a single colony was inoculated in 1 mL YCFA media in a 96 well plate and incubated for 24 hours. Frozen stocks of the 96 well master plates were maintained in glycerol suspension (25% v/v) at -80 °C. Prior to each induction, 96 well plates containing 1 mL YCFA were inoculated from the frozen master plate and grown overnight.

### Bacterial phylogeny and prophage prediction

A set of 40 single copy marker genes were extracted from the 252 bacterial isolates using progenome-classifier[2] and translated into amino acid sequences using SeqKit[3] (v.2). The protein sequences were concatenated and aligned using MAFFT[4] (v7.310) before gaps were trimmed with trimAI[5] (v1.4.1). Maximum-likelihood trees were constructed using RAxML[6] (v8.2.12) PROTGAMMALGF model with 100 bootstraps replicates and visualised in iTOL[7]. Bacterial clusters sharing 99% average nucleotide identity (ANI) were identified using dRep[8] (v.3.0.0) with '-pa 0.9 –sa 0.99' flags. Prophage regions were predicted using Virsorter[9], Vibrant[10] (default settings) and VirFinder[11] (minimum length 5 kb, 0.7 score and p-value 0.05). Completeness was predicted using CheckV[12] and contaminating bacterial regions were removed. Trimmed predictions were located within their cognate bacterial genome and overlapping predictions were merged using R IRanges[13] (v.2.28.0).

**Prophage induction and sequencing**

Prophages were induced by one of two methods. 1) Overnight starter cultures were diluted 1:50 in 1.5 mL standard YCFA media and grown for 5 hours before the addition of Mitomycin C[21] (0.3 or 3 μg/mL, M4287, Sigma-Aldrich) or Hydrogen peroxide[22] (0.5 mM, H1009, Sigma-Aldrich). 2) Starter cultures were diluted 1:50 directly into standard YCFA, YCFA media supplemented with Stevia[23] (3.7 or 37 mg/mL, SweetLeaf, organic stevia leaf extract), carbon depleted media[24] (YCFA media with 50% reduced carbon source) or short-chain fatty acid (SCFA) [25] depleted media (YCFA media without short chain fatty acids). All cultures were then grown for 20-25 hours followed by centrifugation at 4000 xg for 30 minutes and 1 mL supernatants were collected. Supernatants were treated with 10 μg/mL DNase I (DN25, Sigma-Aldrich) and 12 μL RNase A (R6148, Sigma-Aldrich) for 1 hour at 37 °C. Viral particles were precipitated in 7% PEG 0.3 M NaCl overnight at 4 °C, followed by centrifugation at 14,000 xg for 30 minutes after which the pellets were dissolved in 50 μL TE buffer at 4 °C. Next, 20 μL of each sample was mixed with 5 μL loading dye containing 0.8% SDS and 60 mM EDTA and heated at 65 °C for 10 minutes. Samples were loaded on 0.4% agarose gels and run in TAE for 1.5 hours, followed by visualisation of phage sized (~50 kb) DNA bands using Image Studio Lite (LI-COR Biosciences) with a sample to control well signal ratio cut off of 0.03[14]. Samples with suspected viral DNA were treated with 0.5% SDS and 100 μg/mL Proteinase K at 55 °C for 1 hour followed by a 10 minute inactivation at 65 °C. Phenol/Chloroform/isoamyl alcohol (25:24:1) extraction was performed followed by sodium acetate (0.3 M final) and 70% ethanol precipitation with 0.4 mg/mL glycogen overnight at 4 °C. DNA quantity were validated using Qubit (ThermoFisher, U.S) with a minimum of 2 ng/μL required for sequencing. From these a subset of samples were selected for sequencings as follows. First, all samples grown in standard YCFA media or induced with Mitomycin C (except the Fusobacteria isolate which was only sequenced in standard condition) were selected. Second, samples from at least one isolate within a bacterial cluster (99% ANI) grown in the remaining five induction conditions were selected. For 17 clusters, more than one isolate was sequenced in all conditions. Nextera-XT libraries were constructed and sequenced on either Illumina NextSeq2000 or Illumina NextSeq550.

**Regions of interest**

Reads were trimmed using Trimmomatic[15] (v.0.38) (SLIDINGWINDOW:4:25 MINLEN:100) and used to identify induced prophages using two approaches. First, high-quality prophage predictions (>50% completeness) were validated for induction as follows. Read coverage for each library were obtained on their corresponding genome using Bowtie2[16] (v.2.3.5) (default settings). Genome coverage in 100 bp increments were obtained using Samtools[17] (v.1.9) and Deeptools[18] (v.3.1.3) and a custom python script was used to calculate average modified z-score, coverage fold increase and Cohen's D of prophage regions as follows:

[1] $$z\text{-} score_{ave} = mean(0.6745 * (x_p - \tilde{x})/median[x_h - \tilde{x}])$$

where $zs\text{-} core_{ave}$ is the average z-score of the predicted region, $x_p$ is 100 bp coverage increments of the phage region, $x_h$ is 100 bp coverage increments of the host and $\tilde{x}$ is median coverage of the host and

[2] $$Cohen's\ D = \frac{mean(x_h) - mean(x_p)}{\sqrt{\frac{S_h^2 + S_p^2}{2}}}$$

where Cohen's D is the effect size of prophage vs host coverage, $S_h$ and $S_p$ is the standard deviation of the host and phage coverage respectively. Regions with a minimum average modified z-score of 3.5 or an average 2-fold coverage and Cohen's D larger than 0.7 were retained. A custom python script was then used to refine the start stop positions of the prophage regions within each genome, removing flanking 100 bp increments with coverage less than 25% of the mean prophage coverage. In a second approach, regions of increased coverage were identified without previous prophage predictions using hafeZ[19] (v1.0.2) (default settings with -N -S flags). Some of the identified regions of interest were found to be split across several host contigs. To resolve these into full length phage contigs, de novo assembly using MetaViral SPade[20] (default setting) was performed on all libraries. Resulting hafeZ and MetaViral contigs were dereplicated at 99% average nucleotide identity (ANI) over 85% of the alignment fraction using scripts from the CheckV repository and the longest representative within each group were retained for further analysis.

**Identification of induced temperate phages**

Proteins from the resulting contigs of both sets were predicted and annotated using PROKKA[21] (v.1.14.6) (default settings, --hmms) with the PHROGS[22] database. Further, all proteins were scanned against the hmm databases provided by Cenote-Taker2[23] using Hmmer[24] (v.3.3.1) hmmscan (-E 1e-9). To remove potential fragmented protein hits against hallmark genes, a custom phage database was constructed of genomes from Benler *et al.* 2021, Yutin *et al.* 2021, and the INphared database (December 2021 version) dereplicated at 95% ANI over 85% alignment fraction using CheckV scripts[25–27]. The same hmm searches were performed on proteins from the database and half the average length of the middle 80% percentile was calculated as a cut off for each Caudoviricetes hallmark gene (265 AA for terminase large subunit, 245 AA for portal protein and 186 AA for major head protein). To identify any non-Caudoviricetes genomes, HMM searches for Microviridae, Tectaviridae and Inoviridae was performed as follows. Microviridae hallmark VP1 proteins from Wang *et al.* 2019 was made into HMM profiles using MAFFT v.7.310 with standard settings followed by TABAJARA (-t 0.5 -p 50 -w 15 -b 15 -mb 15 -m c -gs 20 -md 3 -cs yes -mb 20)[28,29]. Multiple sequence alignments of the Double Jelly-Roll hallmark protein of Tectaviridae were obtained from Yutin *et al.* 2018 and turned into HMM profiles using HMMER v3.3.1 hmmbuild[30]. These and the Inoviridae protein family HMMs provided by Roux *et al.* 2019 were searched against all proteins using hmmscan (-E 1e-9)[31]. Contigs containing at least one viral hallmark gene were retained.

**Taxonomic annotation and genome analysis**

Viral taxonomy were assigned based on a combination of the protein alignment method previously described in Nayfach *et al.* 2021 against the INphared database and genus level clustering using vContact2[32] against phage genomes in the custom made database used for hallmark gene searches[33]. In cases where the taxonomic assignments from the protein voting and genus level clustering method differed the lowest common classification was assigned. Species level dereplication was performed at 95% ANI over 85% alignment fraction using scripts from the CheckV repository. Diversity generating retroelements (DGRs) were identified using DGRscan[34] with default settings and remote VR regions were identified querying the template repeat using BLASTn[34] (v.2.7.1+) (-dust no -perc_identity 75 -qcov_hsp_perc 50 -ungapped -word_size 4). DGR positive genomes were defined as genomes encoding both a reverse transcriptase gene and containing rerepeat regions.

**Metagenomic read mapping**

The fractional abundance and prevalence of induced prophages within gut viromes were performed as in Benler *et al.* 2021[25]. Briefly, 1232 gut viromes were downloaded from NCBI using bioinfokit (v.2.0.6) and SRA_toolkit (v.2.9), quality filtered using Trimmomatic (v 0.38), decontaminated by removing read aligning to the human genome (GCF_000001405), phage phiX-174 (NC_001422.1) or a collection of cloning vectors (available from ftp://ftp. ncbi.nlm.nih.gov/pub/UniVec/). The decontaminated reads were aligned to induced prophages and the custom phage genome dataset used for taxonomic classification with Bowtie2 (v.2.3.5) (--no-unal --maxins 1000000). Read coverage was obtained using Samtools (v.1.9) and

Deeptools (v.3.1.3) (bamCoverage) and the normalized average fractional abundance was calculated as previously described[35]. A genome was counted as present within a virome if at least 70% of the genome length was covered by reads.

**Cryptic prophage analysis**

Proteins of predicted prophages were predicted using PROKKA v.1.14.6 (default setting, --hmms) and annotated using the PHROG database and gene counts of PHROG categories was obtained for induced, high-quality predictions (>50% completeness) and low-quality predictions (<50% completeness). Percentage gene frequency change of PHROG categories between induced and high completeness (>50% completeness) predictions was calculated for total and presence-absence as follows:

[5] $$Freq.Change\ (\%) = \frac{100*(f_{cry} - f_{in})}{f_{in}}$$

where $f_{cry}$ and $f_{in}$ is the gene frequencies in the high completeness prediction and induced prophage set, respectively. High-quality predictions were aligned to induced prophages using minimap2[36] (v.2.22) (-X -N 50 -p 0.1 -c). Predictions aligning to at least 85% of the active prophages were retained and further filtered to only include hosts that had been sequenced in the same condition(s) as the active prophage. The same search was performed to identify active-active high sequence similarity prophage pairs. The number of HGT and insertion/deletions events between the pairs was calculated for each group using R IRanges (v.2.28.0) and splicejam (v 0.0.77) packages, where an HGT event was defined as a gap within the alignment present in both pairs and insertion/deletion events was defined as a gap present in one of the pairs but not the other. Host ANI of prophage pairs was calculated using fastANI (v.1.33)[37]. dN/dS ratios between prophage pairs was calculated using dRep (compare–SkipMash–S_algorithm goANI) and the dnds_from_drep.py[38] script.

**Differential prophage induction qPCR**

Isolates were streaked onto YCFA plates and grown for 24 hours. Three sperate colonies from each isolate were inoculated into 1 mL YCFA broth and grown overnight. Overnight cultures were diluted 1:50 into 1.5 mL YCFA media and Hydrogen peroxide was added after 5 hours of growth. All cultures were grown for an additional 20 hours and lysates were treated with 2% chloroform and centrifuged for 20 min at 4000 xg at 4 °C and frozen at -80°C until analysis was performed. qPCR was performed in triplicates using SYBR Green I Master Mix (Roche Diagnostics, Mannheim, Germany) with the Roche Lightcycler® 480 system containing 1 µM of each primer, 5 µL of DNA template and 1x SYBR Green I Master Kit, in a final reaction volume of 20 µl. Cycle parameters: initial denaturation at 95 °C for 5 min; followed by 45 cycles of 95 °C for 30 s, 60 °C for 30 s, and 72 °C for 30 s. qPCR primer pairs were custom designed using Primer 3 (https://primer3.org/): Q25 forward 5'-ATCGGTTACGGTCATACGGC-3' and reverse 5'-ATCAAGGGGAAGCGCGTTTA-3', Q41 forward 5'-AAATATGGTGTTCCGCCCGT-3' and reverse 5'-CGTTTCGTTTCCTTGCAGCA-3' and gyrA forward 5'-GGTCATTGTTTCACGTGCCC-3' and reverse 5'-TACCTCACCCACGATTCTGG-3'. *In silico* PCR amplification (http://insilico.ehu.eus/user_seqs/PCR/) did not show cross reactivity of primers to non-cognate prophage and no cross reactivity to the rest of the bacterial genome was found using blastn. Standard curves for primer efficiency analysis were generated via 10-fold dilution in PCR-grade $H_2O$. Samples were diluted 10-fold and qPCRs was performed in triplicates. Efficiency of each primer calculated as in [3] and corrected ΔcT values calculated as in [4]:

[3] $$Efficency = 10^{-1/slope}$$

[4] $$\Delta cT = \frac{EfficiencT_x^{cTx}}{EfficiencT_y^{cTy}}$$

**Long read sequencing**

Isolates were streaked onto YCFA plates and grown for 24 hours. Single colonies were grown overnight in 40 ml of YCFA media, pelleted by centrifugation at 4000 xg for 10 minutes, and washed four times in 1 ml of PBS. DNA was extracted using the Monarch HMW DNA extraction kit (New England Biolabs) following the Gram-positive Bacteria protocol, with modifications. Cells were lysed in 300 $\mu$l of STET buffer (8% sucrose 5% triton x-100 50mM EDTA, 50mM tris Ph 8) containing 10mg/ml lysozyme, 300 $\mu$l of HMW gDNA Tissue Lysis Buffer and 20 $\mu$l of Proteinase K, and incubated at 56 °C for 10 minutes. Lysates were treated with 10 $\mu$l of RNase A at 56 °C for 5 minutes followed by 300 $\mu$l of Protein Separation Solution. Samples were mixed by inversion for 2 minutes then centrifuged at 4 °C for 20 minutes at 16000 x g. Supernatants were collected and 550 $\mu l$ of isopropanol was added to 800 $\mu l$ of supernatant. Samples were inverted for 5 minutes, or until DNA was precipitated, and DNA was pelleted by centrifugation at 4 °C for 10 minutes at 12000 xg. The resulting pellet was washed twice with 500 $\mu l$ of gDNA wash buffer and resuspended in nuclease free water. Library preparation and Oxford Nanopore MinION sequencing was performed using either the Oxford Nanopore ligation sequencing kit (SQK-LSK109) with native barcoding expansion kit (EXP-NBD114) (CC01407, CC01390, CC01401 and CC01405) or the rapid barcoding kit 96 (SQK-RBK110.96, CC01404). Resulting long reads were hybrid assembled with Illumina short reads into closed genomes using dragonfly (v.1.0.14) (CC01407, CC01390, CC01401 and CC01405) or into near complete genome using unicycler[39] (v. 0.4.7) (CC01404) with subsequent scaffolding using RagTag[40] with CC01407 genome as reference.

**Statistical analysis and visualisation**

Significance of PHROG gene category, HGT and insertion/deletion count data calculated with Fischer's exact test and p values adjusted with Hochberg method using R base stats (R version 4.1.3) and rstatix (v.0.7.0) packages. Pearson's correlation test between host ANI and phage pair inducibility as well as Kendall's rank correlation between number prophages within lysogens and prophage inducibility was calculated and plotted using R ggpubr (v.0.4.0) package. Significance of dN/dS data calculated with Wilcoxon test and adjusted by Hochberg method using R ggpubr (v.0.4.0) and rstatix (v.0.7.0) packages. Significance of qPCR fold change between induced prophage in polylysogens calculated with paired t-test (two sided) using rstatix (v.0.7.0), preferential induction calculated with Wilcoxon test and variance of means between isolates calculated with ANOVA using R base stats (R version 4.1.3). Genome maps visualised using R gggenomes (v.0.9.9.9000) package.

**Extended data figures.**

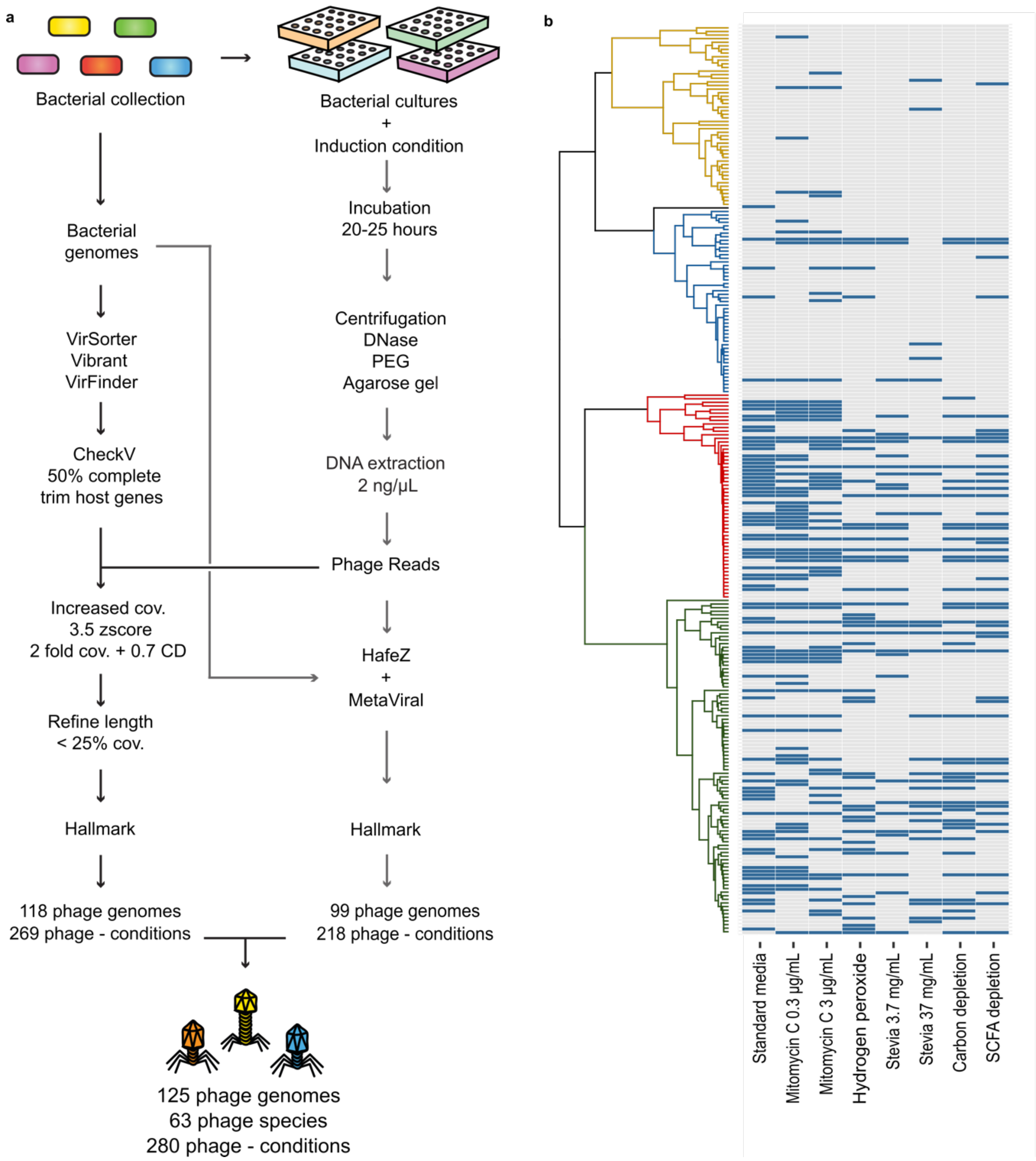


**Extended Data Fig. 1 Methods and sequencing overview. a,** Schematic over the prophage induction and detection protocol. A collection of 252 bacterial gut isolates were grown for 20-25h in either standard media, Mitomycin C (0.3 and 3 μg/mL), Hydrogen peroxide (0.5mM), Stevia (3.7 and 37 μg/mL), carbon depleted or SCFA depleted media (n=2,016). Samples were centrifugated, DNAse treated, PEG concentrated, and phage sized DNA bands were detected on agarose gels. DNA was extracted from samples with phage sized bands (n=861) and samples with DNA concentrations >2 ng/μL were chosen for sequencing based on condition and bacterial cluster (n=431). Reads were aligned to their corresponding bacterial genome and potential induced prophage regions were detected using hafeZ and MetaViral and confirmed as prophages by detection of phage hallmark gene homologues. In tandem, prophage regions were predicted using prophage identification programs and sequences predicted as >50% complete were retained. Reads were aligned to their cognate bacterial genome and predicted prophage regions with >2-fold coverage and Cohen's D >0.7 or mean zscore > 3.5 were retained as potential induced prophage regions. Region were confirmed as viral by detection of hallmark genes. **b,** Phylogenetic tree of the 252 bacterial isolates used for induction assays with heatmap depicting the 431 phage-enriched induction samples chosen for sequencing (sequenced samples shown in blue); Actinomycetota yellow (12/400), Fusobacteriota black (1/8), Bacillota blue (34/408), Pseudomonadota red (174/456) and Bacteroidota green (217/744).

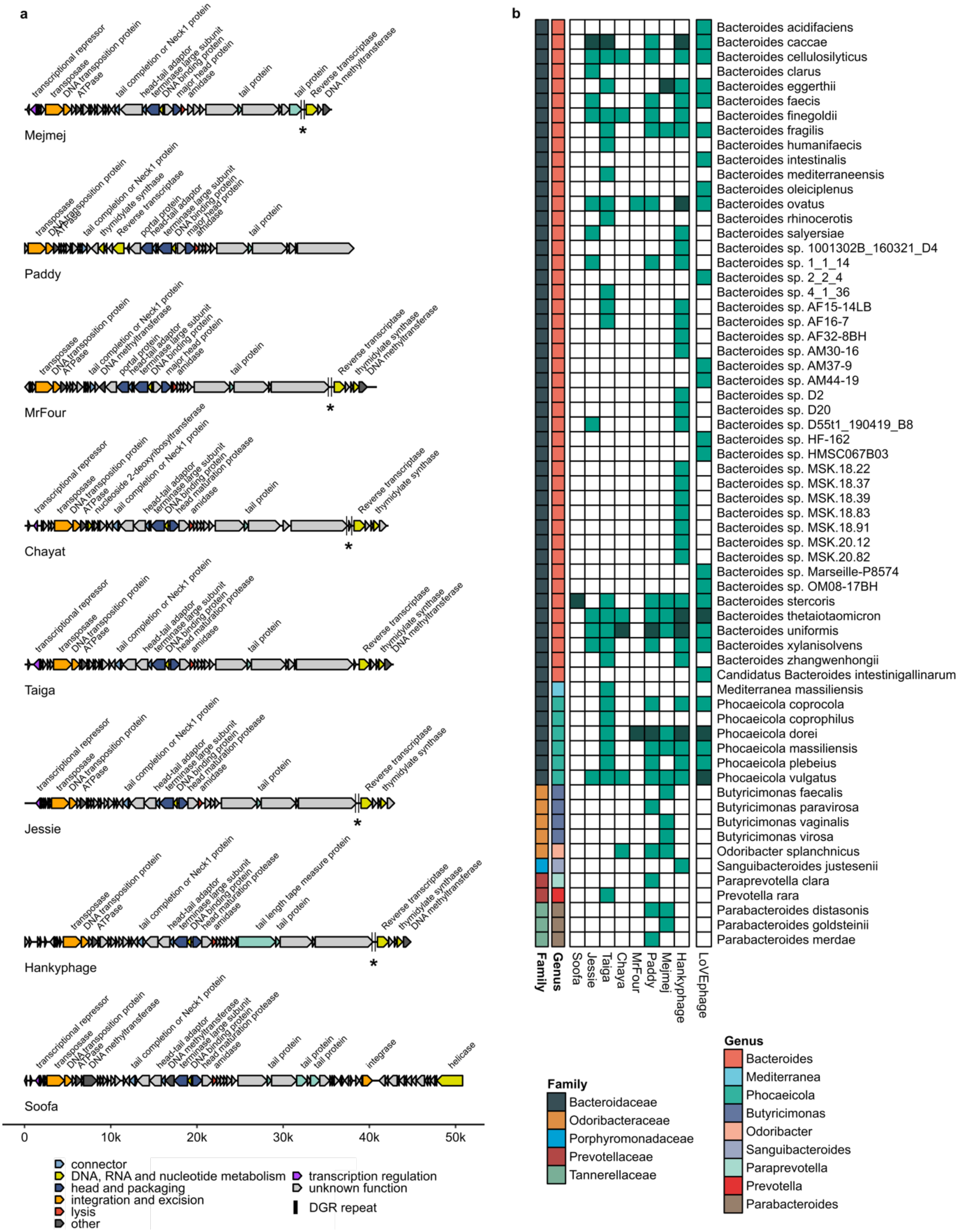


**Extended Data Fig. 2 Hankyvirus genus overview and host range of Hankyvirus and LoVEphage species.** **a**, Annotated genome maps of the eight Hankyvirus genus phages species. Genomes are scaled by length with the ruler displayed at the bottom of the figure. Genes coloured by PHROG categories and unknown genes coloured in grey. DGR template and variable repeats denoted with black lines and highlighted with asterisks. **b,** Host range of Hankyvirus and LoVEphage species induced in this study. Light green denotes hosts found in NCBI RefSeq bacterial genome database; dark green denotes hosts in which the phage was actively induced in this study.

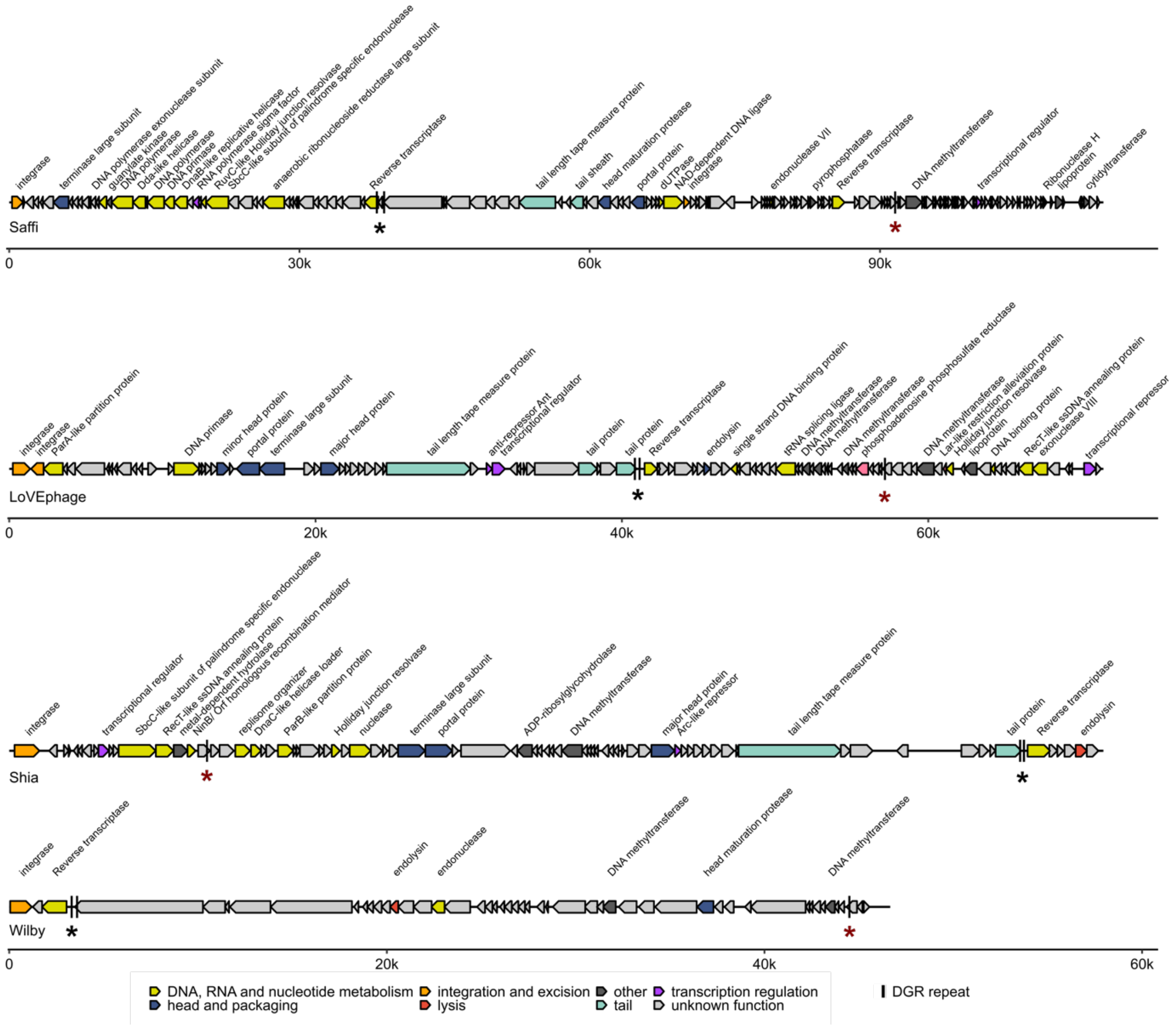


**Extended Data Fig. 3 Genome maps of phage species with double variable repeat DGRs.** Annotated genome maps of three phage species encoding a second variable repeat (VR) distal from the RT cassette. Genes coloured by PHROG categories and unknown genes in grey. DGR template and variable repeats denoted with black lines and highlighted with asterisks; tail targeting region in black and second VR in red.

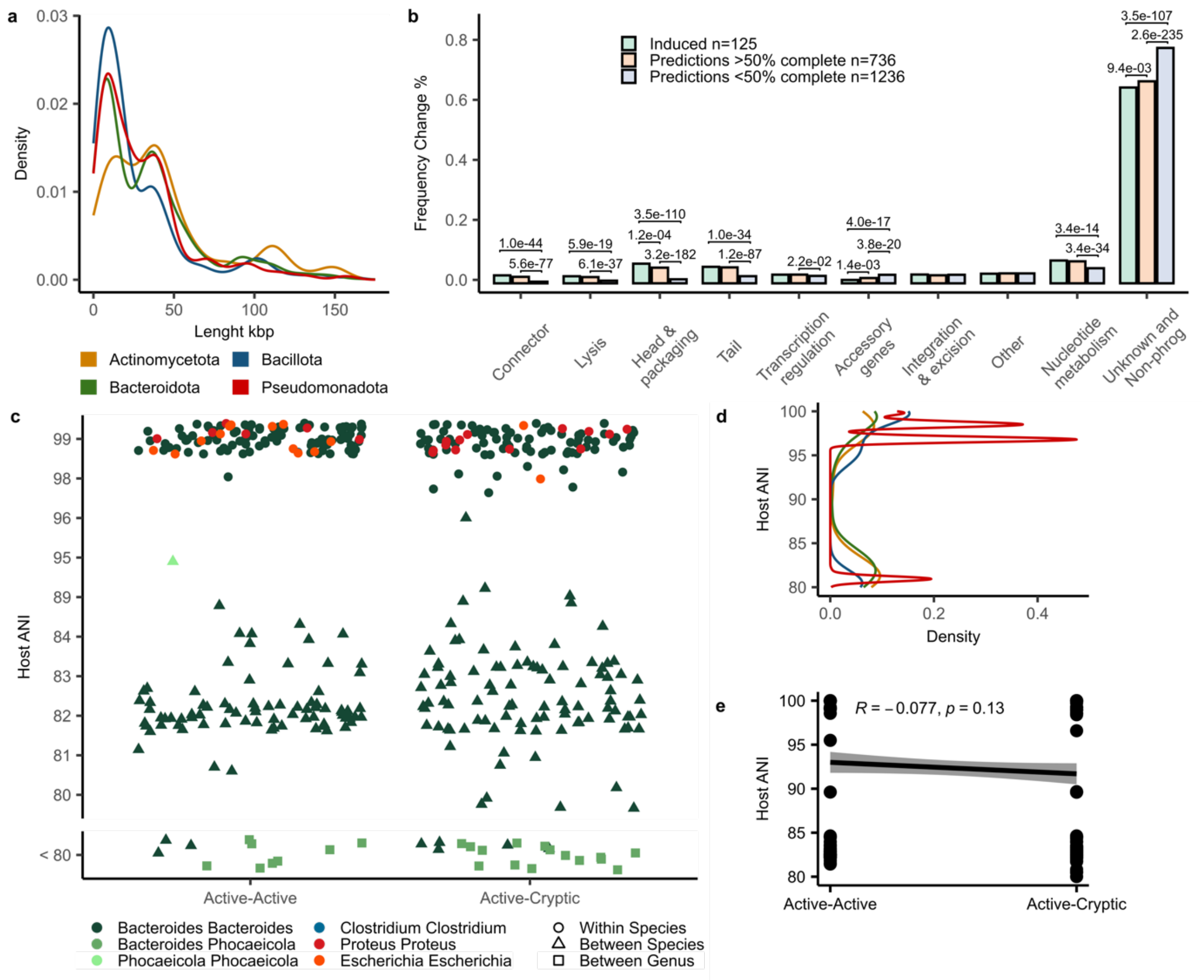


**Extended Data Fig. 4 Comparison of induced versus predicted prophages sequences and host nucleotide identity. a**, Length distribution of all predicted prophages in this study separated by host phyla. A bimodal length distribution was observed for all phyla, with an initial peak at around 8 kb followed by a second peak at around 37 kb. **b**, Frequency of PHROG gene categories across induced (green, n=125), high quality (>50% completeness, orange, n=736) and low quality (<50% completeness, blue, n=1236) predictions. Significant *p* values calculated using Fisher's exact test and adjusted by Hochberg method shown above bars. **c,** Host ANI comparisons of active-active (n=205) and active-cryptic (n=211) prophage homologous pairs. Comparisons coloured by genus and shape based on NCBI taxon. Host pairs with less than 80% similarity grouped separately as too divergent for reliable ANI score. **d**, Density of ANI comparisons of all isolates within the dataset separated by phyla, coloured as in panel a. **c**, Pearson correlation of host ANI between the active-active and active-cryptic prophage pairs. Only ANI values above 80% included for active-active (n=194) and active-cryptic (n=191).
